# Western Ghat Birds Exhibit a Unique Pattern of Seasonal Elevational Shifts and a Combination of Thermal Regime Dependent Shift Strategies

**DOI:** 10.1101/2025.11.05.684726

**Authors:** Ali Khan Faizee, Vijay Ramesh, V. V. Robin

## Abstract

Elevational migration is globally exhibited by montane bird species through elevational range shifts to track seasonal changes in weather patterns and resources. Studies investigating the patterns and mechanisms of such elevational shifts have been limited in spatial scale owing to the difficulty of sampling data across montane gradients. Here, we integrated data from the world’s largest participatory science project (eBird) with systematic surveys to examine patterns and mechanisms that may explain bird elevational range shifts in the Nilgiri Hills of the Western Ghats Biodiversity Hotspot. We calculate the extent of shifts in elevational ranges for birds present year-round in the Nilgiris between the hottest and coldest months and quantify the associations between these shifts and species traits using phylogenetic generalized least squares regressions. We found that 77% (54/70) of the species investigated showed seasonal shifts in their elevational ranges. Birds in the Nilgiris showed nearly equal proportions of downslope (37%) and upslope (33%) winter shifts. All species had significant range overlap between seasons but employed different shifting strategies. Species with narrow thermal regimes predominantly shifted downslope in winter (niche tracking), while those with the widest thermal regimes shifted upslope (niche switching), revealing that species adapt to climatic variability through contrasting thermal strategies. Our results suggest that climatic constraints, specifically thermal regime, could be a major driver of seasonal shifts in elevational ranges of birds in the Nilgiris. Understanding these elevational shift strategies is critical for conservation planning in tropical montane systems facing accelerating climate change.

## 1 Introduction

Elevational migration involves the seasonal movement of a species across elevations annually (Hayes 1995). Several biotic and abiotic hypotheses have been proposed and tested independently to understand the drivers of elevational migration (Vander Pluym and Mason 2024) (See Table 1 for a complete list). Of these hypotheses, climatic constraints, energy efficiency, resource availability, and predation have been purported as key drivers of elevational migration (Barçante et al. 2017; Somveille et al. 2018). The term seasonal elevational shifts is often used in literature instead of elevational migration, because such movements are generally not exhibited by the whole population (Boyle 2017). The most widely tested and applicable, climatic constraints hypothesis, states that the extent of seasonal downslope shifts is a result of their intolerance to climatic seasonality and harshness at higher elevations (Boyle 2011; Cox 1985). Correspondingly, Menon et al. (2023) showed that birds in the Himalayas shift elevations more to remain within narrow thermal regimes (the intra-annual variation in temperature experienced by a species) across seasons. Species also move elevationally as a function of resource availability across seasons. Whereas, the “energy efficiency” hypothesis predicts that a species will be distributed in a way that minimizes costs and maximizes acquisition of energy by targeting areas and elevations with better access to energy supply, as a function of what its competitors are doing (Somveille et al. 2018). Also, different diet guilds like frugivores, insectivores, and nectarivores may move in different vertical directions to avoid resource scarcity (Boyle 2010; Loiselle and Blake 1991). However, research on elevational shift patterns and their drivers has been sporadic across the globe.

**TABLE 1.** Predicted associations between species shifts in elevational ranges and their traits according to the literature and the observed associations between them according to our phylogeny-controlled regression models.

| Hypothesis | Predictor | Upslope shift associations |  | Downslope shift associations |  |
| --- | --- | --- | --- | --- | --- |
|  |  | Predicted | Observed | Predicted | Observed |
| Climatic constraint | Lower temperature limit | – | – | + | + |
|  | Temperature range | + | (+) | – | (–) |
|  | Body size<sup>a</sup> | + | ns | – | ns |
| Food availability | Body size<sup>a</sup> | – | ns | + | ns |
|  | Frugivory/Nectarivory | + | (–) | + | ns |
|  | Generalists | – | ns | – | ns |
|  | Dispersal ability | + | ns | + | ns |
|  | Omnivory | – | ns | – | ns |
|  | Herbivory | – | ns | – | + |
|  | Insectivory | + | ns | + | + |

Note: The associations were examined for upslope and downslope shifts separately. Symbols for the observed associations: + Significantly positive associations under three resampling settings; – Significantly negative associations under three resampling settings; (+) Significantly positive associations under 1–2 resampling settings; (–) Significantly negative associations under 1–2 resampling settings. Abbreviation: ns, no significant associations. <sup>a</sup>The climatic constraint and food availability hypotheses have different predictions on the associations.

Although elevational migration has been documented in at least 10% of the approximately 10,000 bird species worldwide, there is a strong geographic bias in publications focusing on this phenomenon, with a paucity of studies in regions such as the Afrotropical and Indomalayan realms (Barçante et al. 2017). Understanding the factors driving species’ tendencies to shift elevationally is limited, as earlier studies tested these hypotheses on only a handful of species (Boyle 2010, 2011), and more recent comprehensive studies rely solely on participatory science data (Menon et al. 2023; Rueda-Uribe et al. 2024). Among the three non-mutually exclusive hypotheses outlined above, the climatic constraint hypothesis has received the most consistent support, with evidence of thermal regime tracking across multiple regions (Tsai et al. 2021; Menon et al. 2023; Akshay et al. 2024). However, Somveille et al. (2026) recently demonstrated that globally averaged patterns were largely inconsistent with the seasonal climate-tracking prediction and agreed more with the energy efficiency hypothesis. In light of this, we focus primarily on testing the climatic constraint hypothesis. Moreover, elevational shifts may be driven by factors other than, or in addition to, response to climatic fluctuations, for example, dispersal ability and foraging guilds have also been linked to elevational shifts (Sheard et al. 2020; Pageau et al. 2020). A further gap in the literature is the lack of comprehensive comparative investigations, partly because such work requires data volumes that are impractical to obtain through traditional survey methods (Dobkin and Rich 1998).

The use of the world’s largest participatory science project— eBird (Sullivan et al. 2009)—has assisted in identifying widespread and prevalent patterns of elevational shifts for entire bird communities and mountain ranges (Menon et al. 2023; Tsai et al. 2021). Despite its unique advantages and the ability to infer ecological patterns globally, participatory science data are associated with data quality issues, such as spatial clustering, unequal sampling in different seasons and elevations, and undersampling of protected areas (Backstrom et al. 2024; Praveen 2017). Moreover, in topographically complex regions, undersampling (lack of data from several elevational bands due to lack of access and logistical challenges) and oversampling (clustering data from specific elevations alone) can lead to biased inferences (Johnston et al. 2021; Ramesh et al. 2022). This disparity is particularly evident in parts of the understudied Global South, where the elevational gradient is not equally sampled by eBird observers.

Recent methodological developments in ecology have proposed integrating various approaches to biodiversity data collection, such as systematic field surveys, acoustic monitoring, participatory science, and tracking technologies (Miller et al. 2019). Pacifici et al. (2017) developed a framework of integration of semi-structured (participatory science) and structured data (systematic surveys) that addresses the trade-off between data quality and quantity. By using systematic expert surveys to fill the gaps in participatory science data, we can (i) equalize effort across elevational bands, (ii) add sampling effort in inaccessible terrain and protected areas, and (iii) reduce biases in species detection effort. Some studies have demonstrated that combining data from participatory science with systematic surveys improves the ecological inference and predictive ability of distribution models (Miller et al. 2019; Robinson et al. 2020; Zhao et al. 2024). Such an integrative approach is critical in tropical mountains where sufficient studies and data are not available to quantify patterns and shifts in elevational ranges of birds. We use this approach of combining systematic surveys and participatory science to examine seasonal elevational shifts in birds at the scale of a mountain system in the Western Ghats of India.

The Western Ghats are a mountain range running from north to south along the west coast of India. The Western Ghats possess a 2600 m elevational gradient and encompass an array of weather profiles that vary throughout the year, thus potentially affecting the intra-annual variation in temperature, food resources, and precipitation. The Nilgiris of the southern Western Ghats provide an interesting opportunity to examine the patterns of elevational shifts. As the Nilgiris exhibit a variety of thermal niches, they offer the opportunity to test the support for the climatic constraints hypothesis, and also uncover the associations of other species traits with elevational shifts using its high resident bird diversity. Across the eastern slopes of the Nilgiris, we quantify seasonal shifts in elevational migration and infer the potential explanatory factors of these movements at a fine scale.

In this study, for 70 species of birds present year-round in the Nilgiris, we used a combination of systematic surveys and participatory science to:

1. determine the extent and patterns of shifts in elevational ranges,
2. test whether thermal regime drives shifts in elevational ranges, and
3. determine the associations between these shifts and species traits like diet, dispersal ability, and body mass.

Menon et al. (2023) showed that Himalayan birds shift downslope in the winters to track their thermal regimes. The Nilgiris are relatively aseasonal and elevationally restricted compared to the Himalayas. Thus, we expect a much smaller magnitude of elevational shift in birds of the Nilgiris; however, it is difficult to predict the pattern of their associations with thermal regimes, and they may show more than one of four potential patterns (Figure 1): (A) Species with the largest shifts have the narrowest thermal regimes, implying a negative relationship between shift magnitude and thermal regime width (niche tracking); (B) species with the largest shifts have the widest thermal regimes (niche switching); (C) species with narrow thermal regimes show intermediate shifts, while those with wide thermal regimes show either the smallest or largest shift; (D) elevational shifts may be unrelated to thermal regime altogether, instead being shaped by other life-history traits such as dispersal ability, body size, or dietary guild.

**FIGURE 1.**
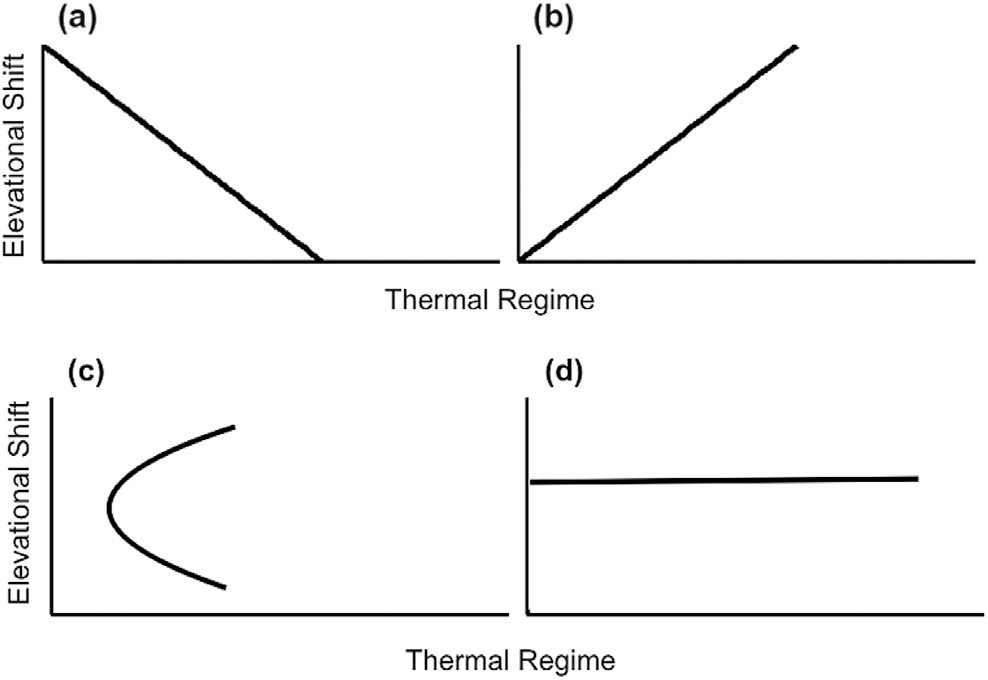
Conceptual plot for pattern of elevational shifts where (a) taxa with the largest shift have the narrowest thermal regime (niche tracking), (b) taxa with the largest shift have the widest thermal regime (niche switching), (c) taxa with the smallest and largest shifts have the widest thermal regimes, (d) no relationship between elevational shift and thermal regime.

## 2 Methods

### 2.1 Study Site

The Western Ghats are a mountain range running for 1600 km along the west coast of India. These mountains encompass a range of latitudes and a wide range of habitats. The Nilgiris is a mountain range in the southern Western Ghats with a range of habitat types (Figure S1), from scrub forests (below 600 m a.s.l.) to tropical montane cloud forests (above 1400 m a.s.l., locally known as *Shola*) and mid-elevation broadleaf forests (between 700 m and 1400 m a.s.l.) (Gadgil and Meher-homji 1986; Pascal et al. 2004). The 2600 m elevational gradient of the Nilgiris experiences a unique thermal regime across elevations (Figure 2a, Figure S2). The Nilgiris support a high diversity of about 287 resident bird species (Viswanathan et al. 2024), of which 16 are globally threatened (IUCN 2024). We focus on the eastern slope of the Nilgiris due to the extensive participatory science observations and a road network covering the entire elevational gradient (Figure 2b).

**FIGURE 2.**
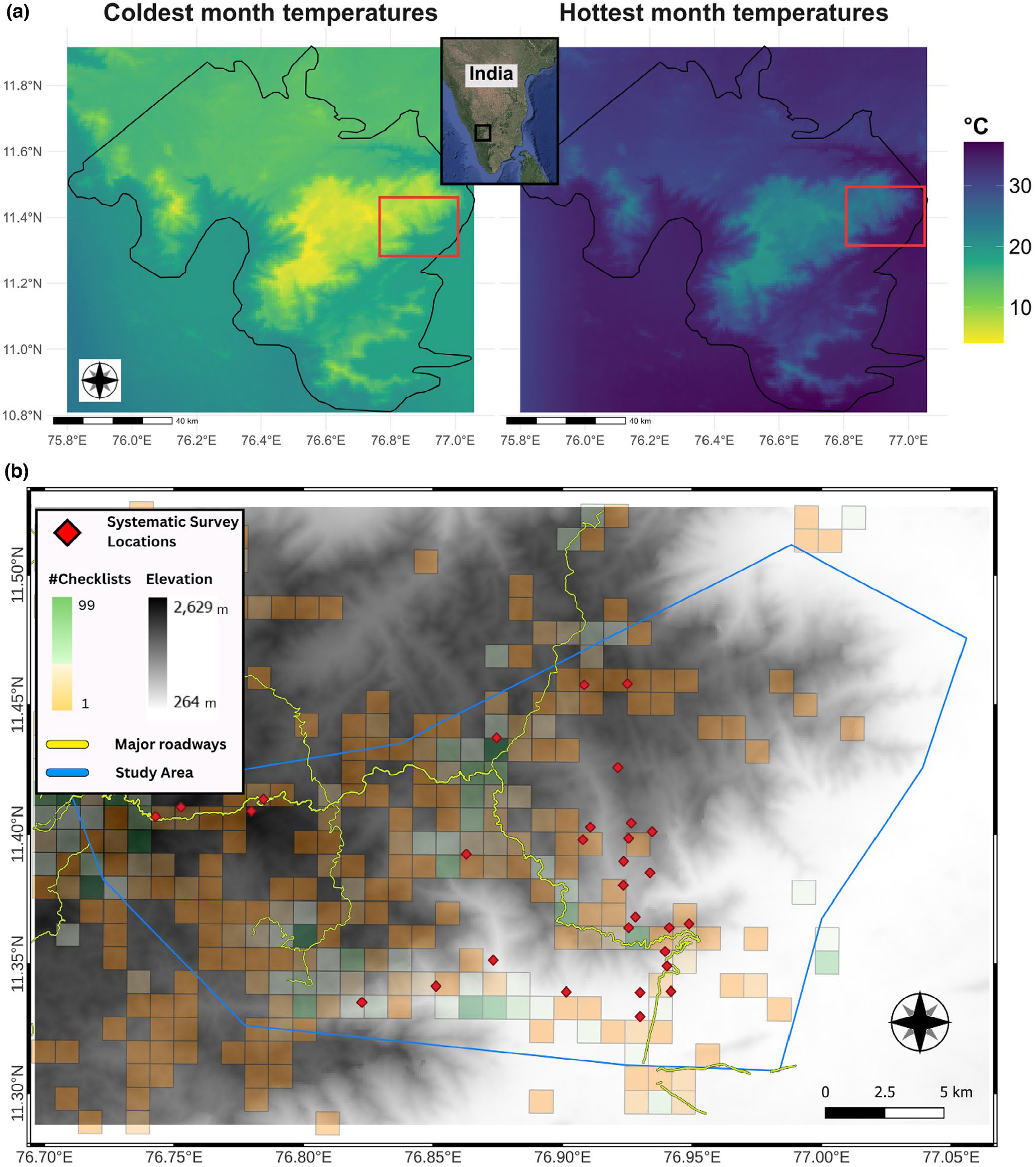
The Nilgiris mountain range, bounded by a black line in (a), is a group of mountains in the Western Ghats of India, and the red square denotes our study area. (a) Left: Mean daily maximum air temperature of the warmest month (May); right: Mean daily minimum air temperature of the coldest month (January). The black outline marks the Nilgiris Mountains of the Western Ghats. (b) Map showing the distribution of eBird sampling effort (checklists) along the elevational gradient of eastern Nilgiris; the orange to green color gradient of the 1 km^2^ squares denotes the number of checklists within the square, and red diamonds indicate locations of our systematic surveys.

### 2.2 Curating eBird Data

We used eBird (Sullivan et al. 2009), a global, crowd-sourced platform collecting bird observations, to extract species occurrence information across the eastern slope of the Nilgiris (Figure 2b). Data on eBird consist of observations uploaded by users and stored as checklists. Each checklist contains information on bird species presence, number of individuals, observers, time spent observing birds, etc. We initially downloaded all eBird data from 2013 to 2022 for our study area (31,524 checklists). This dataset resulted in checklist-level information for 935 unique localities (unique GPS coordinates of checklist locations) spanning elevations ranging from 100 m to 2600 m above sea level. We determined the hottest and coldest quarters for the eastern Nilgiris mountain range using bioclimatic data at 1 km resolution from CHELSA (Karger et al. 2021). We then divided the eBird dataset into two subsets based on the month of the sampling event. The summer dataset consisted of data from the hottest quarter in the study site (i.e., April to June). Similarly, the winter dataset included data from the coldest quarter (i.e., November to January).

From the filtered eBird dataset, we retained unique, complete checklists with the following attributes in our dataset: distance traveled less than 2.5 km, checklist duration not more than 2 h, and checklists with fewer than ten observers, following Menon et al. (2023) to reduce spatial uncertainty and data quality issues. Further, we filtered out nonresident species in the Nilgiris by selecting only those species listed as resident (birds that do not show any latitudinal migration) in both the State of India’s Birds report (SoIB 2023) and the Kerala Bird Atlas (Praveen et al. 2022). We removed species that are predominantly aerial, prone to misidentification, or nocturnal, including Accipitridae, Apodidae, Hirundinidae, Caprimulgidae, Falconidae, and Strigiformes. See Table S1 for a complete list of species included and excluded in the analysis.

### 2.3 Generating and Integrating Data From Systematic Surveys

Since participatory data are opportunistic, sampling varies across certain regions and seasons (Backstrom et al. 2024), introducing biases and gaps. We explored the number of checklists per 100 km^2^ for various elevational bins (200 m, 400 m, 500 m). We found that 400 m elevational bins provide the most evenly distributed checklist density across elevational bands (Figure S3). We then conducted systematic surveys across locations with limited eBird data to fill these gaps and improve downstream analysis. We found the distribution of eBird checklists was clustered near major highways and in certain elevational bands (Figure 2b). From these grids, we selected three locations at least 400 m apart from each other, and more than 200 m away from any road (to avoid disturbances) in each 400 m elevation band as spatial replicates for our systematic field surveys.

In each chosen location, we carried out a 20-min variable-distance point count (Dawson et al. 1995), recording all visual and aural observations of bird species. We sampled 30 localities (Figure 2b) spanning 264 m to 2200 m in elevation four times, at least thrice in the morning and at least once in the evening, between December 2023 and May 2024. We generated a combined dataset by collating bird species occurrences from systematic surveys and eBird. We reorganized our systematic point-count surveys to resemble the structure of the checklist data from eBird. Each point count was treated as a stationary count (distance traveled = 0), and a unique sampling identifier was generated by concatenating the plot ID and date. After removing the non-relevant variables (like last edited date, subspecies, country, IBA code, media, comments, etc.), the two datasets were identical in structure, which allowed us to combine them. The resulting dataset is referred to as the combined dataset throughout the text. A graphical pipeline of data cleaning, integration, and downstream analysis processes is shown in Figure 3.

**FIGURE 3.**
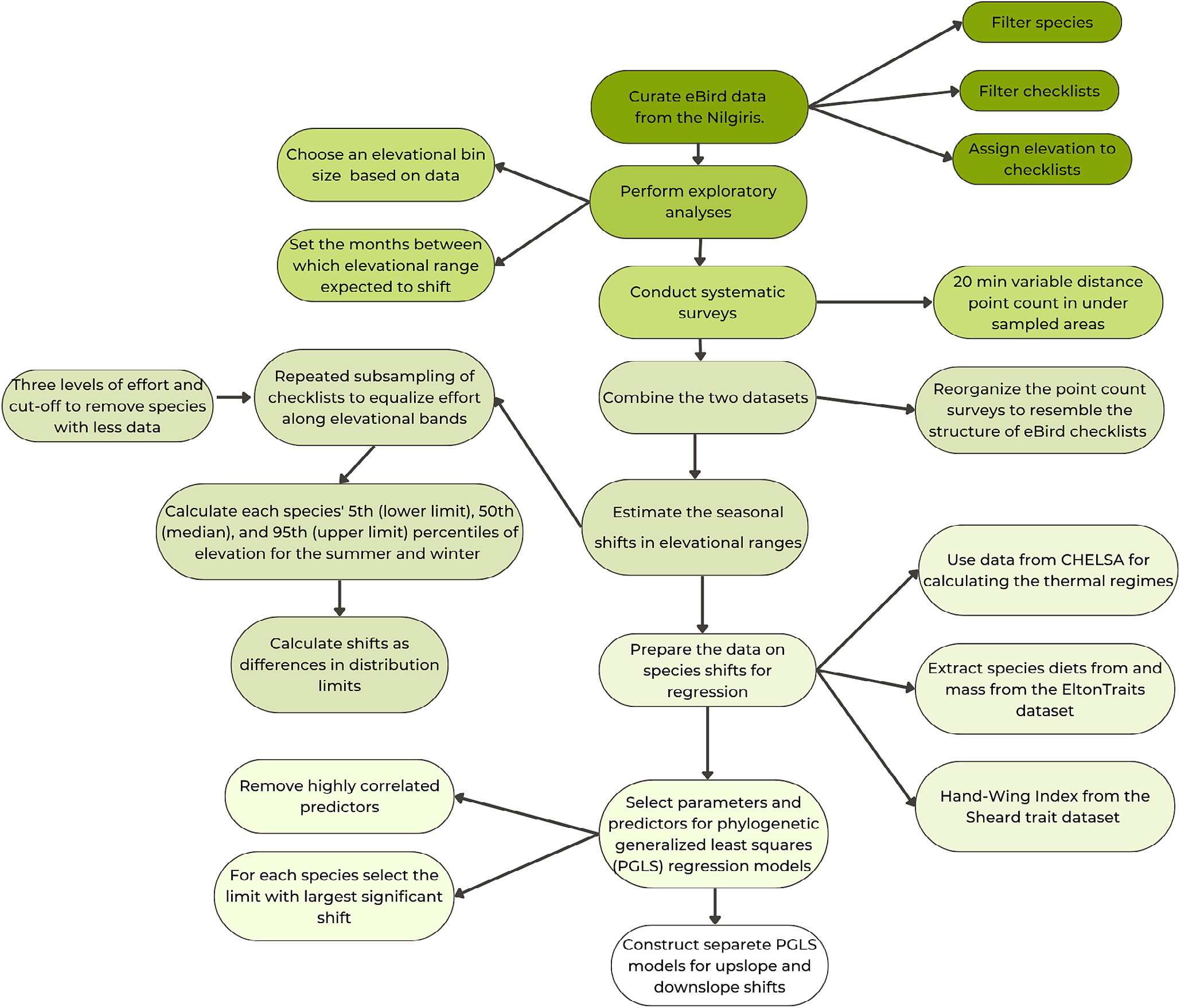
A graphical pipeline of data cleaning, integration, and downstream analyses process.

### 2.4 Estimating the Seasonal Shifts in Elevational Ranges

We extracted the elevation for each sampling locality of the combined data using the SRTM Digital Elevation Model (NASA JPL 2013). We calculated the average elevation in a 100 m radius to account for detection distance and uncertainty in GPS locations. Although we conducted systematic surveys along elevational bands, the data thus generated were not independently comparable to the eBird dataset. To account for sampling bias arising from uneven survey effort across elevations, we employed a resampling approach described by Tsai et al. (2021). The resampling approach generates subsets of data with a controlled effort (number of checklists) from each elevational band by repeatedly subsampling (with replacement) a fraction of checklists from the whole dataset on a predefined effort parameter (1st, 2nd, and 3rd quartiles of checklists distributed across elevation bands). We achieved equal sampling effort across elevation band-season combinations by subsampling checklists with replacement 1000 times.

This procedure equalizes the density of checklists across the different elevation bands, because the elevation bands with larger areas will have more checklists by default, which will get sampled more often. To account for fluctuations brought about by subsampling, we performed it with three different levels of effort: the first, second, and third quartiles (500 checklists, 1673 checklists, and 4095 checklists, respectively) of the total sampling events among elevation bands and seasons. To avoid bias in elevation ranges for uncommon species and species with low detection rates, we only included species with more than 15 occurrences in all elevational bands each season for further analysis. This number was based on the threshold used by Menon et al. (2023) and Tsai et al. (2021), who used a threshold of 30 records for carrying out similar analyses in the whole of the Himalayas and the country of Taiwan, respectively, and compared that number (30) with a higher threshold of 60 and 100 records and found not much difference in the ranges. Given our study site, the Nilgiris has an area less than half of these studies, we chose to use 15 as a threshold to not lose out on many species. Since 15 was randomly selected, we also repeated the analyses for 30 occurrences. Using the resampled datasets, we calculated each species’ 5th, 50th, and 95th percentiles of elevation for the summer and winter seasons separately, representing the lower limit, center of distribution (median), and upper limit, respectively. The seasonal shifts were then calculated as differences in distribution limits (upper limit, lower limit, or median) in two seasons (summer vs. winter). We classified shifts in species distribution parameters (lower limit, median, and upper limit) into downslope shifts if the 95% confidence interval of the difference in their distribution limits (summer minus winter) was above zero, upslope shifts if this interval was below zero, and no shift if it contained zero. In total, 70 species had at least 15 records in both seasons in at least one resampled sampling event set and were included in the following analyses.

### 2.5 Determining Potential Predictors of Elevational Shifts

To test the climatic constraint hypothesis, we checked whether the lowest (lower temperature limit), highest (higher temperature limit) temperatures a species experiences in a region, and thermal regime (the difference between the highest and lowest temperatures in an area) are associated with shifts in elevational ranges. Climatological data comprising maximum and minimum temperatures for January (coldest month in the Nilgiris) and May (hottest month in the Nilgiris) were sourced from CHELSA at a 1 km resolution (Karger et al. 2017). These data were used to derive mean minimum January and mean maximum May temperatures for each zone per 100 m elevational band. Each species’ thermal regime was quantified as the difference between the mean maximum May temperature at its summer elevation and the mean minimum January temperature at its wintering elevation, with separate calculations performed for the center, lower, and upper limits of its elevational ranges. For each species, we define the mean maximum June temperature at its summer elevation and the mean minimum January temperature at its wintering elevation as its higher and lower temperature limits. The higher and lower temperature limits for the species had a large positive correlation (Table S2), so we only used the species’ lower temperature limit in further analyses.

We examined the association between species’ diets and shifts in elevational ranges. We extracted species diet preferences (frugivory, nectarivory, omnivory, herbivory, and insectivory) from the EltonTraits dataset (Wilman et al. 2014) and used omnivory as the reference category for the regression analysis. Frugivory and nectarivory were clumped together because there is high synchrony in fruiting and flowering, respectively, and significant overlap between flowering and fruiting in the Nilgiris (Jeevith and Kunhikannan 2019; Murali and Sukumar 1994; Krishnan 2004). Consequently, frugivores and nectarivores tracking these resources should exhibit similar movement patterns. Research across the southern Western Ghats supports this, showing strong phenological overlap in fruiting and flowering peaks in habitats ranging from montane evergreen forests (peaking in March) to the tropical dry forests of Mudumalai and the understory in Kalakad–Mundanthurai Tiger Reserve (Krishnan 2004).

We calculated the dietary diversity of a species by calculating the Shannon-Wiener diversity index from the proportions of food types in the diet listed in the EltonTraits dataset, which we used as a proxy for a generalist diet (Fontaine et al. 2008). We used body mass on a base 10 logarithm scale (to reduce the huge variation between species) to find whether body size correlates with shifts in elevational ranges. Finally, we used the hand-wing index from the Sheard trait dataset (Sheard et al. 2020) as a proxy for dispersal ability (Arango et al. 2022) to find whether it is correlated with shifts in elevational ranges.

We separated upslope and downslope movements because their ecological drivers differ, leading to different expected relationships between species traits and elevational shifts (Table 1; Vander Pluym and Mason 2024). This distinction is particularly important in our study, since we expect a high frequency of upslope shifts in the tropical Nilgiris, a pattern that contrasts with the downslope-dominated shifts reported from the subtropical and temperate Himalayas (Menon et al. 2023). A species was assessed to have an upslope shift if any one parameter (lower limit, median, or upper limit) shifted upslope without any other parameter shifting in the opposite direction. Similarly, downslope shifts were also assessed as a congruous shift.

We constructed phylogenetic generalized least squares (PGLS) regression models (Martins and Hansen 1997) to determine the associations between shifts in either of the three measures of elevational distribution (lower limit, median, and upper limit) and the traits mentioned above. Using our combined dataset, we calculated the shifts in elevational range for each species (lower limit, median, and upper limit). Further, resident species whose subpopulations are involved in latitudinal migrations were also removed from the PGLS analyses, because these range shifts may be due to the migratory populations moving into the Nilgiris in winters.

Before running the PGLS models, we removed predictors that strongly correlated with each other, meaning the mean highest temperature in the hottest quarter, which was correlated with the mean lowest temperature in the coldest quarter, was removed. Further, for the upslope shift model, the species’ parameter (one from the lower limit, median, or upper limit) that showed the largest significant upslope shift (95% confidence interval of the shift below zero) was selected for the PGLS regression. Similarly, for the downslope shift model, the species’ parameter (one from the lower limit, median, or upper limit) that showed the largest significant downslope shift (95% confidence interval of the shift above zero) was selected. Splitting the species movements into upslope and downslope reduces the number of species and thereby the statistical power of the analysis because the difference between the number of predictors and instances of significant shifts (also referred to as cases in regression terminology) is reduced. Thus, we checked if there were significant cases in the models and found that there were at least twice as many cases as predictors in each model (Tables S3.1 and S3.2), which retains considerable statistical power (Mundry 2014).

We iterated over the different levels of sampling effort (first, second, and third quartiles) and occurrence threshold datasets (15 and 30 detections) to build models. A phylogenetic consensus tree was obtained with the least-squares edge lengths method from the 1000 trees (downloaded from the Bird Tree of Life; Jetz et al. 2012) and was pruned to the species present in the analysis, according to the requirements of the PGLS algorithm. We assumed that a Brownian Motion evolution model could explain the error structure in the residuals, and these errors correlate with the species’ phylogenetic closeness. All analyses were carried out in R version 4.3.3 with the help of the phytools (Revell 2024), nlme ver. 3.1 (Pinheiro et al. 2018), and ape (Paradis and Schliep 2019) packages.

## 3 Results

We recorded 1685 observations of 91 resident species from our systematic field surveys. We recorded data from 132 species across 19,234 checklists for the same region. Using the combined dataset, we retained 70 species having enough occurrences (at least 15 in each season), which was an improvement from the 55 species retained when using the eBird dataset alone. We found that 54 (77%) species seasonally shifted at least one of their upper, lower elevational limits or median across seasons (Figure 4, Table S4), and 16 species did not shift their ranges. The elevational range reduced for 24 species from summer to winter, while it expanded for 28. Thus, a pattern of partial elevational range shift was prevalent. The magnitude and prevalence of upslope and downslope shifts were not significantly different. We found a median downslope shift of 153 m, 282 m, and 287 m, and a median upslope shift of 129 m, 369 m, and 225 m in the lower limit, median, and upper limit, respectively.

**FIGURE 4.**
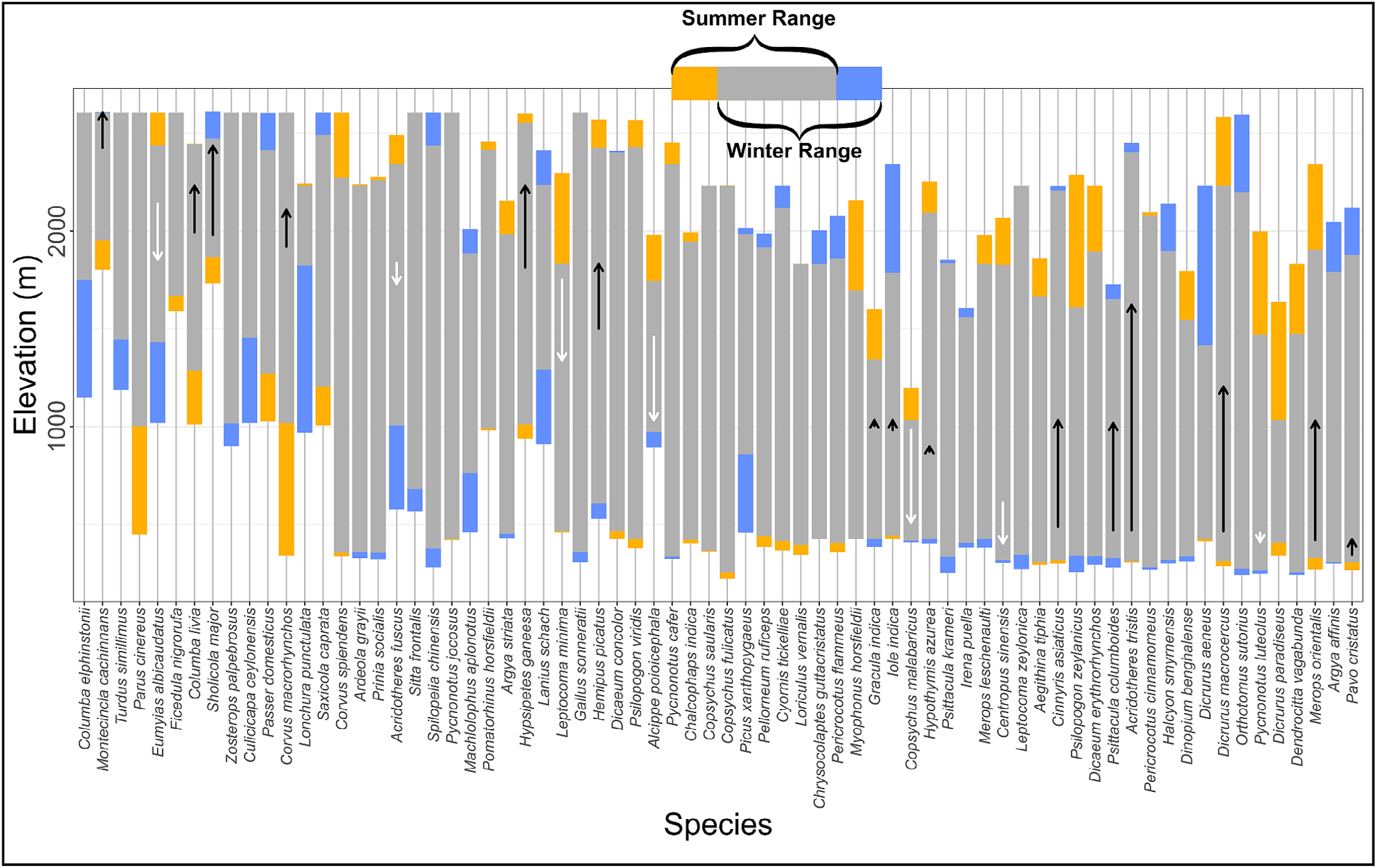
Elevational range comparisons in the Nilgiris (threshold = 15) between summer and winter. Yellow and blue bars represent the elevational ranges estimated during summer and winter. The gray represents the overlap in range between these two seasons, and arrows indicate a significant difference in the median (the length of the arrow is proportional to the difference).

We found that 26 of these 70 species had a significant downslope shift (95% confidence interval of shift estimate completely below zero) and no upslope shift in either the lower limit, upper limit, or median, indicating downslope movements during winter (Table S4). Similarly, 23 species had a significant upslope shift (95% confidence interval of shift estimate completely above zero) during winter and no downslope shift in the lower limit, upper limit, or median, indicating upslope movements. We found that species shifted their lower limits more often than their upper ones. Specifically, 43 species shifted their lower limits, 22 shifted their median, 15 shifted their upper limits, and 22 species shifted more than one of their limits. This pattern matched the trend in temperature range (the difference between the highest and lowest temperatures experienced in that elevational band), which was greater at lower elevations (Figure S2). Also, 7, 14, and 7 species shifted their lower limit, median, and upper limit, respectively, more than 200 m. The birds that show the largest shifts in their lower limit, median, and upper limit of distribution are *Lonchura punctulata* (855 m down in winter), *Acridotheres tristis* (1121 m up in summer), and *Psilopogon zeylanicus* (625 m down in winter), respectively. *Culicicapa ceylonensis* and *Turdus simillimus* show an expansion in their winter range (i.e., the upper limit moved higher and the lower limit moved down).

The PGLS model that examined downslope shifts had a residual standard error = 70.19 and 18 error degrees of freedom. We discovered consistent and large negative associations between shifts in elevational ranges and the range of temperatures experienced by the species (relative effect = −142 m ± 7.10 (SE, standard error), *t*_28_ = −36.69, *p* < 0.05). In addition, significant positive associations were detected between the lowest temperature experienced by a species (lower temperature limit) and downslope shifts (relative effect = 15 m ± 4.8 (SE), *t*_28_ = 4.64, *p* < 0.05) (Figure 5a). Across diet guilds, with omnivory as a reference, we did not find any significant associations with downslope shifts (Figure 5a). The PGLS model that examined species showing upslope shifts had a residual standard error = 54.12 and 9 error degrees of freedom. We recovered a positive association between the range of temperatures experienced by a species and upslope shifts in elevational ranges (relative effect = 134 m ± 7.00 (SE), *t*_20_ = 27.46, *p* < 0.05). We also found a significant negative effect of the lowest temperature experienced by a species (lower temperature limit) on upslope shifts (relative effect = −15.1 m ± 2.6 (SE), *t*_20_ = −8.16, *p* < 0.05). Only in one of the upslope shift models did we find a significant negative association of species eating more fruits and nectar with upslope shifts, compared to omnivores (Figure 5, relative effect = −76.1 m ± 23.4 (SE), *t*_20_ = −3.25, *p* < 0.05). Species dietary diversity, species dispersal ability (hand-wing index), and size (body mass) did not seem to have significant effects on the extent of elevational shift (*p* > 0.05, Table S5) in any of the models. Notably, downslope shifting birds moved within significantly narrower thermal regimes, while upslope shifting birds showed larger shifts for wider thermal regimes (Figure S4).

**FIGURE 5.**
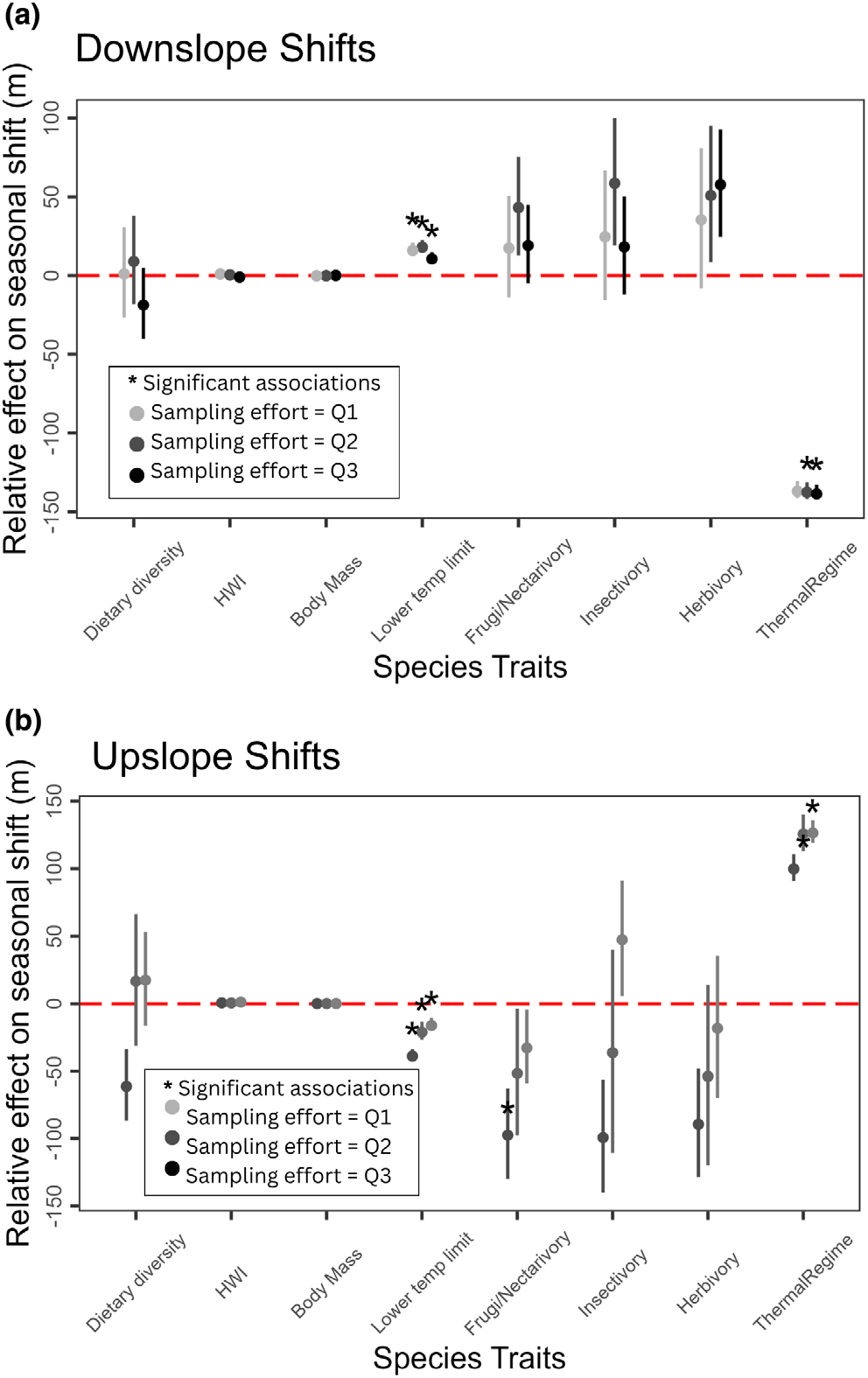
Associations between birds’ traits. (a) Downslope shifts in their elevational ranges, (b) upslope shifts in their elevational ranges, in the Nilgiris. Asterisks denote significant effect sizes (*p* < 0.05). Dots are the model estimates, and error bars represent standard errors for PGLS models. The dot color indicates the sampling effort (light gray to black: low to high). Estimated from the eBird + systematic surveys dataset. For the trait dietary preference (Frugi/Nectrivory, Insectivory, Herbivory), the reference category is omnivory, and all the effects shown in the figure are relative to the reference category.

## 4 Discussion

In this study, we characterized the patterns and associations between seasonal shifts in elevational ranges and various traits of 70 resident bird species in the eastern Nilgiris of the Western Ghats biodiversity hotspot. We show that the traditional survey methods (Dobkin and Rich 1998), which are financially and logistically complicated at large spatial scales, can be integrated with eBird data to infer ecological patterns even for topographically complex areas. Our analysis suggests that 77% of the selected species in the resident bird community exhibit seasonal shifts in their elevational ranges. However, this pattern was not recovered uniformly across all species, as overlaps in summer and winter ranges were common, and most species had specific ways of shifting their ranges between seasons. This lack of nonoverlap in seasonal elevational ranges in the Nilgiris can be attributed to the smaller elevational gradient (0–2600 m), limited temperature seasonality, and the tropical climate promoting year-round availability of resources, as compared to other montane ecosystems such as the Himalayas or the Andes (Imfeld et al. 2021; Karuppusamy 2023; Yadav et al. 2021). We found that species exhibiting seasonal elevational shifts in the Nilgiris use two contrasting thermal regime dependent shift strategies. Species with narrow thermal regimes predominantly shifted downslope in winter, adopting the niche tracking strategy, while those with the widest thermal regimes shifted upslope, switching their niche.

Although shifts of small magnitude were common, we found that a few species shifted either the upper limit, the lower limit, or the median of their distribution more than 200 m (*n* = 24). Among these relatively large shifts, a more than 200 m change in the median of the distribution was most common (*n* = 14), followed by the lower and upper limits (*n* = 7 for both). This indicates possible shifts in relative abundance along the elevational gradient rather than the population moving as a whole. We found two patterns for upslope shifts: (i) generalist human-associated species (like *Passer domesticus, Columba livia*, and *Pavo cristatus*) making large upslope shifts, which could be to exploit less competition at higher elevations during winters; (ii) locally adapted species (like *Chrysocolaptes socialis* and *Iole indica*) may be moving upslope to breed in extensive tree forests of mid and high elevations during retreating monsoons in the eastern slope of the Nilgiris (December to February). We also note that the high-elevation endemic species (shola species, e.g., *Montecincla cachinnans, Ficedula nigrorufa*) do not show large shifts in elevational ranges in the eastern Nilgiris, except for the significant downslope shift for *Sholicola major*.

We find subtle differences in the elevational shift patterns and their associations with species traits inferred in this study, focused on the Nilgiris and another one that targeted the entire Western Ghats (Akshay et al. 2024), and speculate that these differences could arise due to different geographic scales and the pipelines of analyzing participatory science data. The proportion of species in the eastern Nilgiris showing elevational shifts in our analysis (77%) is notably higher than a recently reported estimate (35%) from the Western Ghats (Akshay et al. 2024), but comparable to estimates in temperate Himalayas (65%) and other subtropical regions (70%) (Menon et al. 2023; Tsai et al. 2021). When shifts in elevational ranges are calculated at the scale of a large latitudinal gradient (e.g., the Western Ghats), mountain-specific patterns in elevational shifts may go undetected due to averaging across multiple mountain ranges and movements across mountains. Moreover, the variations of area in elevational bands, sample size for each bird species, and uneven distribution of biases can contribute to the differences between these studies.

We found that the thermal regime and the species’ lower temperature limit best explained the patterns of elevational shifts in the Nilgiris. In agreement with studies in various latitudinal zones, our results support the climatic constraints hypothesis (Hahn et al. 2004; Menon et al. 2023; Tsai et al. 2021). We found that species with a narrower thermal regime have a higher tendency to shift their distribution downslope in winters (niche tracking), while species with the largest upslope shifts had the widest thermal regime (niche switching). The high elevations (> 1400 m) experience the lowest temperatures and lower seasonality in the Nilgiris, and we found many mid-elevation species (species with medians of distribution around 1000 m) to be using these elevations only in the summer. Taken together with the overall positive association of shifts and species’ lower temperature limit, it appears that birds shifting downslope in the Nilgiris are tracking their thermal regime. We speculate that the reasons for downslope movements in summer could vary among species: for example, to exploit fruiting, seasonal insect abundances, or to search for breeding sites. Moreover, the opposite association with upslope shifts (implying birds that often shift their ranges upslope experience lower minimum temperatures) suggests that some species might be adapting the niche switching strategy and shifting upslope to exploit seasonal resources in certain elevational bands. Recent studies have found empirical evidence of bird elevational migration consistent with the energy efficiency hypothesis, including upslope movement during the colder season as a distributional strategy that minimizes energetic costs in the context of resource competition (Somveille et al. 2026). Our findings about the upslope shifts in the Nilgiris are consistent with this hypothesis. Moreover, given that the low elevations in the Nilgiris (0 to 1000 m) experience the highest temperatures and temperature seasonality, the finding that several birds in these elevations do not show large elevational shifts in their lower distributional limits suggests that the lower temperature limit may determine their distribution more strongly. Also, the moderate negative correlation (Pearson’s correlation of −0.4) between thermal regime and the medians of the species distributions implies that species have wider thermal regimes in the lower elevations and narrower ones in the higher elevations.

Our finding that diet guild had no major significant associations with shifts in elevational ranges, combined with the conflicting evidence from other similar studies (Menon et al. 2023; Tsai et al. 2021), suggests that the abundance of food resources along this elevational gradient may not vary drastically between seasons, or that birds may have variable food preferences. A detailed investigation of seasonal resource availability, interspecific competition, foraging opportunities, and food preferences along the elevational gradient of the Nilgiris remains a promising avenue for future research.

While our approach can be relied on to study patterns of elevational shifts, there are a few limitations. We acknowledge that biases in eBird data can persist even after strict filtering (Backstrom et al. 2024), and participatory science datasets at small scales are prone to fluctuations due to outliers (Weisshaupt et al. 2021). We tried to address these limitations using complete checklists, repeated resampling, excluding the data with species identification uncertainties, and filling spatial sampling gaps with systematic surveys. We also recognize that multiple abiotic and biotic processes often act in conjunction to determine migration patterns (Janzen 1987); however, testing such intricate processes was beyond the scope of our study. We also note that our tests on foraging and thermal regime associations with elevational shifts are from secondary data, and direct measurements may provide detailed insights.

In conclusion, we inferred the patterns of seasonal shifts in elevational ranges of a subset of resident birds in the Nilgiris and uncovered associations of these shifts with species traits in this tropical mountain range. While doing so with a combination of participatory science data and systematic surveys, we demonstrate the advantage of taking an integrative approach to fill gaps and infer ecological patterns even for topographically complex, undersampled areas. We found that resident birds in the Nilgiris exhibit a unique pattern of seasonal elevational shifts and a combination of thermal regime dependent shift strategies. Our results suggest that climatic constraints, specifically thermal regime, could be a major driver of seasonal shifts in elevational ranges of birds in the Nilgiris. Additionally, beyond intraspecific dynamics, interspecific competition for energy may also play a meaningful role (Somveille et al. 2018 and Somveille et al. 2026); the Nilgiris serve as winter grounds for 82 migratory species, whose seasonal influx could intensify resource pressure on residents at particular elevational bands, potentially driving elevational shifts. Yet given our limited understanding of how migratory and resident species interact across shared ranges (Cohen and Satterfield 2020), the energy efficiency hypothesis warrants more rigorous investigation. Resolving these ecological interactions has direct implications for conservation planning in tropical montane systems increasingly exposed to the pressures of climate change. Conservation must prioritize maintaining intact elevational gradients and connectivity in protected areas to facilitate species movements (Elsen et al. 2018), as habitat fragmentation limits species’ ability to track suitable climates and adapt to warming (Neate-Clegg et al. 2021).

## Acknowledgments

We thank the Ministry of Environment, Forest and Climate Change (MoEFCC), Government of India (through their Long-Term Ecological Observatories Programme), and the National Geographic Society for financial support of this study. We extend our thanks to Ashwin Warudkar, Vinay KL, Chiti Arvind, Dr. Jobin Varughese, Naman Goyal, and Archita Sharma for their insightful feedback and constructive criticism, which have played a crucial role in shaping the direction of this project. We thank the Tamil Nadu Forest Department for providing us with permits and support to carry out the field component for this research. Furthermore, we appreciate M. Mubeen, Kovaithambi, Subramanian, Chandrasekar Das, Anita Varghese, and the Keystone Foundation, whose assistance has been indispensable in completing the fieldwork. We also thank Dr. Ramana Athreya, Harikrishnan CP, Dr. Geetha Ramaswami, Meera MR, Aravind P.S., Zeba Madani, and Dev Bagdi for their timely guidance and advice. Lastly, we would like to sincerely thank the anonymous reviewers, whose constructive feedback and thoughtful suggestions greatly enhanced the quality of this manuscript. Additionally, we acknowledge the Indian Institute of Science Education and Research Tirupati, for their support in this research.

## Funding

This work was supported by the National Geographic Society (30422123) and the Ministry of Environment, Forest and Climate Change, Government of India (13008/72/2019-CC).

## Ethics Statement

No specific ethical clearances were sought for this project, as it did not involve working with people or invasive research about plants/animals. The legal permit to carry-out bird surveys was provided by the Tamil Nadu Forest Department- Permission No. 71/2023.

## Conflicts of Interest

The authors declare no conflicts of interest.

## Data Availability Statement

The data that support the findings of this study are available in figshare at https://doi.org/10.6084/m9.figshare.31609885, reference number 31609885. These data were derived from the following resources available in the public domain: eBird—http://ebird.org/data/download and CHELSA-bioclim—https://www.chelsa-climate.org/datasets/chelsa_bioclim. The R scripts used to analyze these data are archived at https://github.com/faizee-ali/ElevationalMigration.git.

## Supporting Information

Additional supporting information can be found online in the Supporting Information section. **Figure S1:** (a) the red outline encompasses the Nilgiris Mountain Range, courtesy of Google Earth 10.85 (2015); (b) the area in each 50 m elevation band in the Nilgiris Mountain Range in km^2^ calculated from the SRTM digital elevation model (OpenTopography, 2013). **Figure S2:** (a) Range of temperatures (b) mean of minimum temperature in December along the elevational gradient in the Nilgiris, derived from averaging climatic data from the bio7 (Temperature Annual Range = bio5—bio6) and bio6 (Min Temperature of Coldest Month) layers in the CHELSA dataset (Karger et al., 2017) for each 100-m elevational band, respectively. **Figure S3:** Sampling effort (checklists per 100 km2) for each 400 m elevational band in the eastern slope of Nilgiris for (a) summer (March–May) and (b) winter (Nov–Jan). Sampling events (checklists) along the y-axis and elevation along the x-axis. **Figure S4:** Relation between thermal regime and elevational shifts for (a) downslope shifts and (b) upslope shifts. **Table S1:** List of bird species of the Western Ghats, with is_selected column indicating the species we selected for this study can be found here: https://doi.org/10.6084/m9.figshare.31609885. **Table S2:** Table of Pearson’s correlation values between the covariates used in PGLS regression. deltaElev is Elevation, HWI is Hand-Wing Index, minTemp_Dec_mean is the mean lowest temperature for each elevation band, maxTemp_May_ mean is the mean highest temperature for each elevation band, diet_s_i is the Shannon–Wiener diversity index for the diet, and mass is the log of mass of the species. **Table S3.1:** Data used for running the phylogenetic generalized least squares regression on downslope movements. Can be found here: https://doi.org/10.6084/m9.figshare.31609885. **Table S3.2:** Data used for running the phylogenetic generalized least squares regression on upslope movements. Can be found here: https://doi.org/10.6084/m9.figshare.31609885. **Table S4:** List of species with more than 15 occurrences in both seasons, and the estimated shifts in their lower limits, medians, and upper limits, in the Nilgiris mountain range can be found here: https://doi.org/10.6084/m9.figshare.31609885. **Table S5.1:** Model output from the phylogenetic generalized least squares regression exploring the relationship between seasonal elevational downslope shift and diet, dispersal ability (HWI, Hand-wing index), dietary diversity (diet_s_i, Shannon Index of food types), body mass (mass), and thermal regime breadth can be found here: https://doi.org/10.6084/m9.figshare.31609885. **Table S5.2:** Model output from the phylogenetic generalized least squares regression exploring the relationship between seasonal elevational upslope shift and diet, dispersal ability (HWI, Hand-wing index), dietary diversity (diet_s_i, Shannon Index of food types), body mass (mass), and thermal regime breadth can be found here: https://doi.org/10.6084/m9.figshare.31609885.

## Notes

https://doi.org/10.6084/m9.figshare.30061018.v1

### Competing Interest Statement

The authors have declared no competing interest.

### Summary of Updates

proofread and accepted manuscript from Biotropica

